# Transcriptomic Analysis Reveals MSTN Mutations and Mechanisms of Muscle Hypertrophy in a New Guinea Pig Breed

**DOI:** 10.1101/2024.12.09.627469

**Authors:** Sheyla Carmen, Lilia Chauca, Claudia Yalta, Enrique Alvarado, Edwin Mellisho

**Author notes:** Corresponding author: (EM).

## Abstract

The guinea pig (*Cavia porcellus*), a species native to Peru, is valued for its meat production, where muscle development is essential for productive efficiency. The new Kuri breed, obtained through selective breeding and genetic selection, has shown a phenotype with more developed musculature compared to native guinea pigs. In this study, we conducted a comparative transcriptomic analysis between Native and Kuri breed guinea pig, complemented by histological analysis of the semitendinosus muscle to investigate the underlying mechanisms responsible for the differences in muscle morphology. Histological analysis revealed a significant increase in muscle fiber area in the Kuri breed compared to the native guinea pigs. At the molecular level, key mutations were identified in the *MSTN* gene, including variants in the 3’ UTR region and a frameshift mutation, which alter the gene’s inhibitory function on muscle growth. Additionally, differences were observed in the expression of pathways related to muscle degradation, energy metabolism, and angiogenesis, which explain the greater muscle hypertrophy in the Kuri breed. These findings provide a first understanding of the genetic mechanisms responsible for muscle hypertrophy in the Kuri breed and suggest candidate genes for improving meat quality through molecular genetic breeding programs in guinea pigs.

**AUTHOR SUMMARY:** The guinea pig in Peru is valued for its meat production. The new Kuri breed, obtained through selective breeding and genetic selection has shown a phenotype with more developed musculature compared to other breeders. In our study, comparative transcriptomic analysis between Native and Kuri breed guinea pig, complemented by histological analysis of the semitendinosus muscle to investigate the underlying mechanisms responsible for the differences in muscle morphology. Histological analysis showed significant increase in muscle fiber area in the Kuri breed compared to the native. Furthermore, it is the first time that we identify in guinea pig, key mutations in the MSTN gene, which alter the gene’s inhibitory function on muscle growth. The impact of this finding will allow us to plan genetic improvement strategies in this species that promote muscle hypertrophy, paving the way for future research and its potential impact on guinea pig production.

## INTRODUCTION

The guinea pig (*Cavia porcellus*), a native animal originating from the Andes of Latin America [1,2], is highly valued in Peru for its cultural importance and its growing role in meat production for human nutrition [3,4]. Peru leads the commercial breeding of guinea pigs, with a population that has successfully expanded across various ecological regions of the country [5], which has allowed for significant growth of this industry in rural areas.

In this specie, the muscle development directly affects productive efficiency; however, the lack of standardization in variables such as nutrition, management, genotype, age, and sex at the time of slaughter causes carcass weights to vary considerably, ranging from 237 g to 893 g, with carcass yield fluctuating between 34.8% and 73.4% [6]. Among the carcass, the most important cuts are the leg and ribs, which represent between 35% and 40%, followed by the shoulder (14-17%) and neck (6.5-10.8%) [6,7].

Differences in the chemical composition of the meat from various guinea pig breeds, such as Peru, Andina, Inti, Inka, Merino, and Criollo, have been reported [8]. These variations, along with differences in muscle development based on age and sex [7], emphasize the need to investigate the genetic bases influencing these traits to improve guinea pig production.

The Kuri (new breed, released in 2021) after a long process of phenotypic and genealogical selection and directed crossbreeding between breeds such as Peru, Inti, and Andina, has stood out for its higher meat yield and reproductive capacity [9].

Compared to other breeds, the Kuri has shown a phenotype of greater musculature and superior prolificacy, being 13.5% more prolific than the Peru breed and with a body weight 20% greater than the Andina and Inti breeds. These characteristics make it an ideal breed for commercial meat production, generating growing interest in studying the genetic factors underlying its muscle development.

Among the key genes related to muscle development, myostatin (MSTN) stands out as a negative regulator of muscle growth that limits the proliferation and size of muscle fibers [10]. The structure of MSTN is highly conserved [11], underscoring its importance in the regulation of muscle growth across various species. The function of MSTN as a growth inhibitor has been widely demonstrated in several animal models. For example, mice with mutations that inactivate MSTN show a significant increase in muscle mass compared to wild-type animals, highlighting the critical role of this gene in regulating muscle development [12]

This pattern is repeated in larger species, where mutations or MSTN knockout have resulted in the “double-muscle” phenotype, characterized by a notable increase in muscle mass, such as in pigs [13,14], rabbits [15–17], goats [15, 18, 19], buffalo [20], catfish [21], sheep [19, 22]), and chickens [23]. The study of MSTN function in guinea pigs is especially relevant, as native breeds exhibit more limited muscle growth compared to kuri breed, offering a unique opportunity to investigate the molecular mechanisms behind muscle hypertrophy.

In this study, we conducted a comparative transcriptomic analysis between Native and Kuri breed guinea pigs to identify differences in gene expression that could explain the muscle development observed in the Kuri breed. Additionally, we complemented this analysis with histological studies of the semitendinosus muscle. These combined approaches allowed us to delve into the genetic and biologic mechanisms responsible for muscle growth in this new guinea pig breed, providing a foundation for future research on the genetic improvement of this species.

## RESULTS

### Muscle histology

Increasing the production of meat for animal consumption is one of the ultimate goals of sustainable animal husbandry [34]. Genetic selection to improve body weight and the selection of genes that control the growth of muscle mass are considered practical ways to improve meat production [35,36]

The fiber type composition of the semitendinosus muscles was analyzed by cross section (Fig 1; Fig S1). The results revealed that on average the semitendinosus muscle fiber area was 927.07 µm² for the native guinea pigs and 1760.21 µm² for the kuri breed, highlighting the differences in fiber size between the two groups. These findings offer significant insights into the muscular characteristics of these types of guinea pigs and their potential implications for their physiology and overall performance.

**Fig 1.**
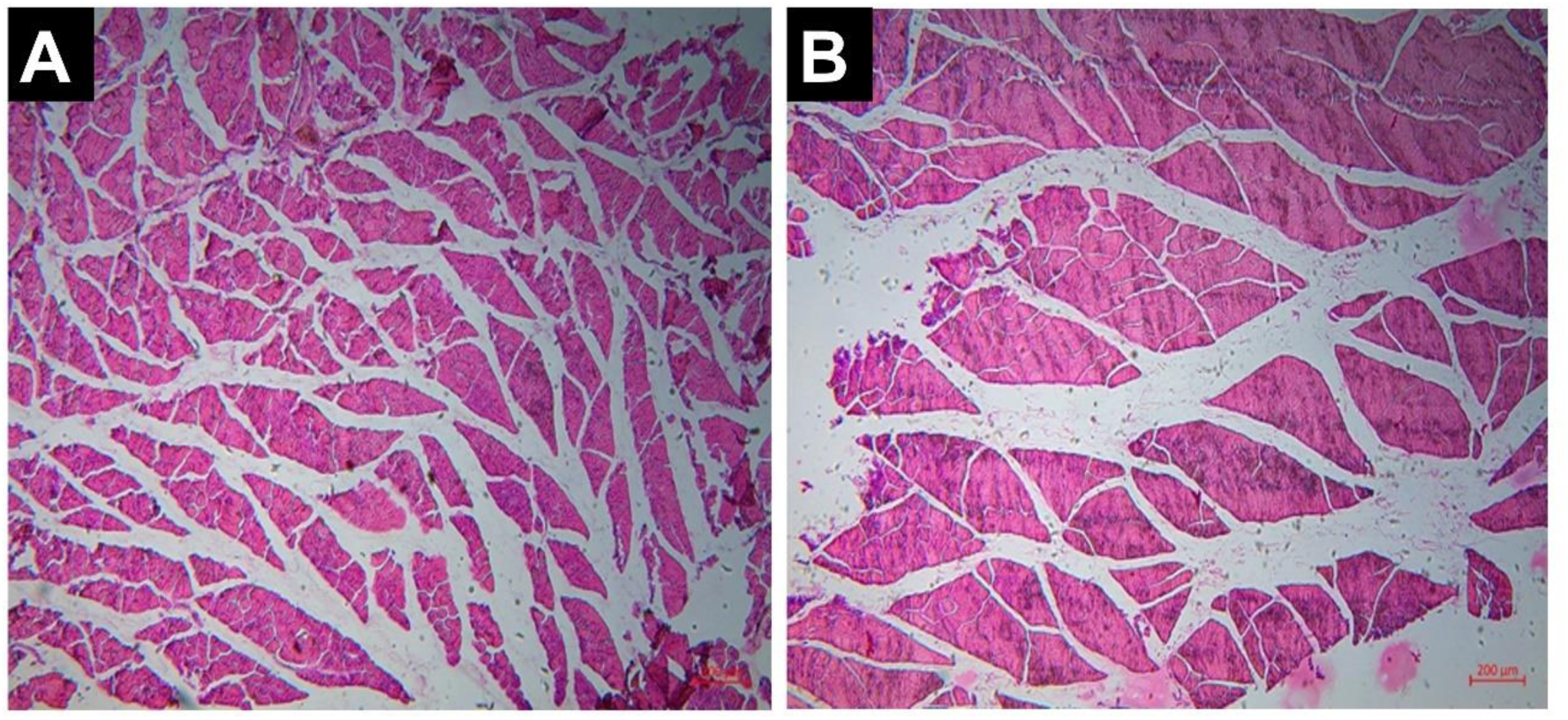
Histological section of the guinea pig muscle stained with Hematoxylin-eosin (40x). A) Native and B) Kuri breed.

### RNA-Seq Data Analysis

In total, 6 cDNA libraries composed of two groups (native and Kuri breed guinea pig) with three replicates for each group were sequenced. This generated a pool of 263 million 150 bp reads, with a range of 40 to 48 million reads per sample. A constant error rate of 0.03 was obtained in all the samples, and in all of them, the Q20 quality values exceeded 97%, while the Q30 quality values exceeded 93%. In general, good quality results were found in all the analyzed samples (Table 1 and 2). Also, we observed that ∼90.8% of the reads aligned to the reference genome. In addition, we obtained percentages of ∼82.1% for the reads that aligned in a single location, exceeding the threshold of 80% which would indicate it’s a good library [37]. Additionally, we found a lower percentage of reads that aligned to multiple locations (∼8.8%) and thus, reduce the error due to multi-mapped reads that can negatively impact the calculated gene expression levels [38]. A total of 16146 genes were detected in guinea pig semitendinosus muscle tissues, of which 15130 genes exhibited shared expression in all six samples (Fig 2B).

**Fig 2.**
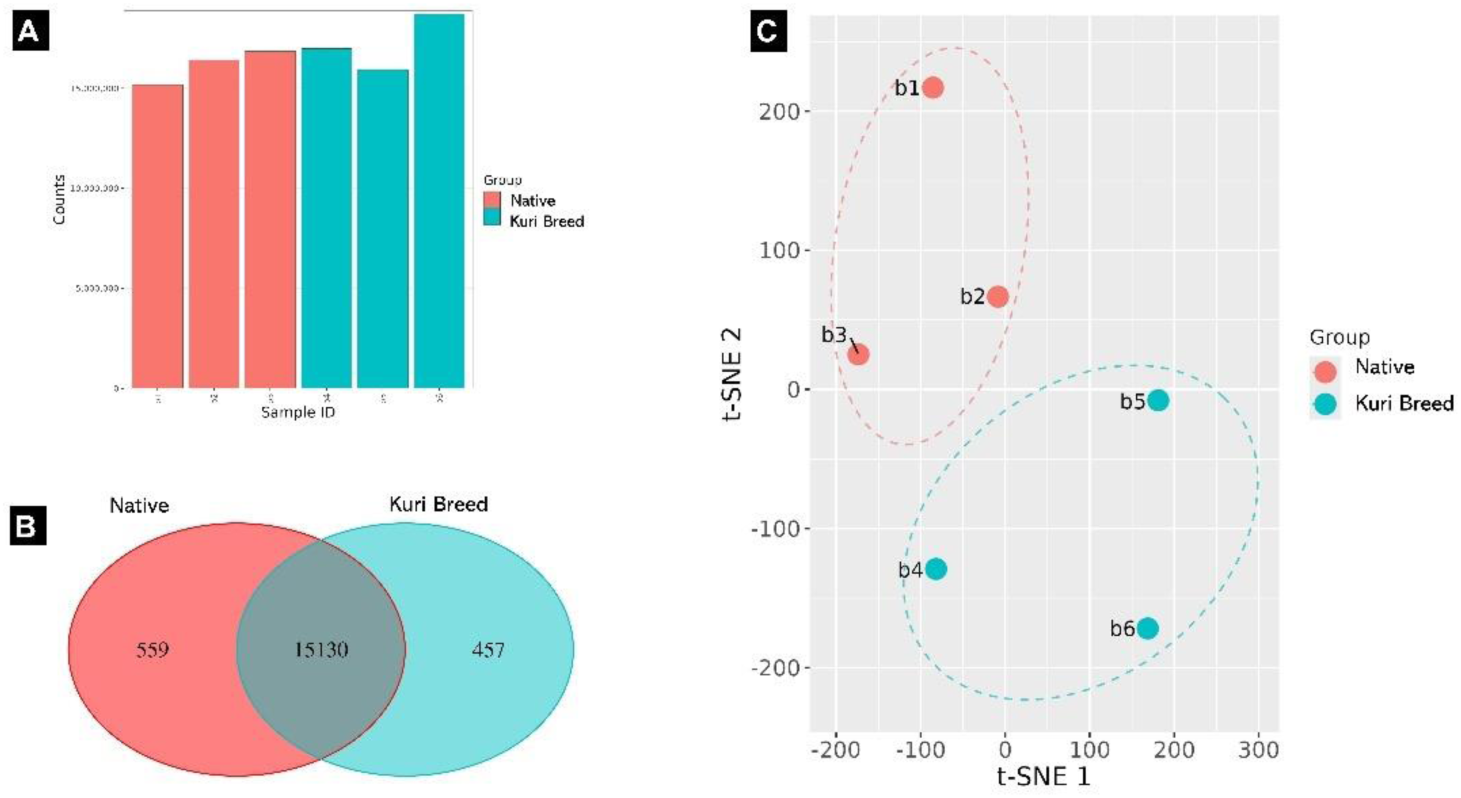
Distribution of global gene expression of muscle tissue of Native and Kuri breed guinea pig. (A) Distribution of clean bases of sequencing reads in global gene expression of muscle tissue of Native (red) and Kuri breed (blue) guinea pigs. (B) Venn diagram depicting genes that are uniquely expressed found in global gene expression of muscle tissue of Native (red) and Kuri breed (blue) guinea pigs, (C) t-SNE in global gene expression of muscle tissue of Native (red) and kuri breed (blue) guinea pigs.

**Table 1.** Data quality control summary.

| Group | Sample | Raw reads | Clean reads | Error Rate | Q20 (%) | Q30 (%) | GC content (%) |
| --- | --- | --- | --- | --- | --- | --- | --- |
| Native | B1 | 41393400 | 38463038 | 0.03 | 97.76 | 93.81 | 51.75 |
|  | B2 | 43666320 | 40972776 | 0.03 | 97.73 | 93.7 | 51.33 |
|  | B3 | 44370240 | 42146352 | 0.03 | 97.96 | 94.22 | 50.67 |
| Kuri breed | B4 | 44330086 | 42541668 | 0.03 | 97.71 | 93.62 | 51.19 |
|  | B5 | 40604424 | 39783540 | 0.03 | 97.87 | 93.95 | 50.98 |
|  | B6 | 48754532 | 47658652 | 0.03 | 97.96 | 94.11 | 49.49 |

**Table 2.** RNA-Seq mapped events.

| Group | Sample | Total reads | Total mapped |  | Uniquely mapped |  | Multiple mapped |  |
| --- | --- | --- | --- | --- | --- | --- | --- | --- |
|  |  |  | Reads | (%) | Reads | (%) | Reads | (%) |
| Native | B1 | 38463038 | 35019707 | 91.05 | 31461622 | 81.80 | 3558085 | 9.25 |
|  | B2 | 40972776 | 37472103 | 91.46 | 34129014 | 83.30 | 3343089 | 8.16 |
|  | B3 | 42146352 | 38071119 | 90.33 | 34044748 | 80.78 | 4026371 | 9.55 |
| Kuri breed | B4 | 42541668 | 38402516 | 90.27 | 34882658 | 82.00 | 3519858 | 8.27 |
|  | B5 | 39783540 | 36568649 | 91.92 | 33617205 | 84.50 | 2951444 | 7.42 |
|  | B6 | 47658652 | 43322328 | 90.90 | 38458427 | 80.70 | 4863901 | 10.21 |

### Differential Expression Analysis

In the differential expression analysis, we identified a total of 209 differentially expressed genes in the semitendinosus muscle of Native and Kuri breed guinea pig (DESeq2 padj ≤ 0.05, |log2FoldChange| ≥ 1.0). Of these, 44 genes were upregulated, while 165 genes were downregulated, representing more than three times the number of downregulated genes compared to upregulated ones (Table S1 and S2). These results suggest that the main phenotypic differences native and Kuri breed guinea pig are largely due to the silencing of genes and pathways rather than the activation of new molecular routes.

Among the most notable upregulated genes were *MIOX* and *TTC39C*, while genes such as *GADD45G*, *FBXO32*, *DDIT4*, *TNFRSF12A*, and *MAP3K8* were downregulated in the kuri breed guinea pigs (Fig 3A). An important observation was that the *MSTN* gene showed slightly elevated expression in the kuri breed, although it narrowly failed to surpass the threshold for upregulation (|log2FoldChange| = 0.935).

**Fig 3.**
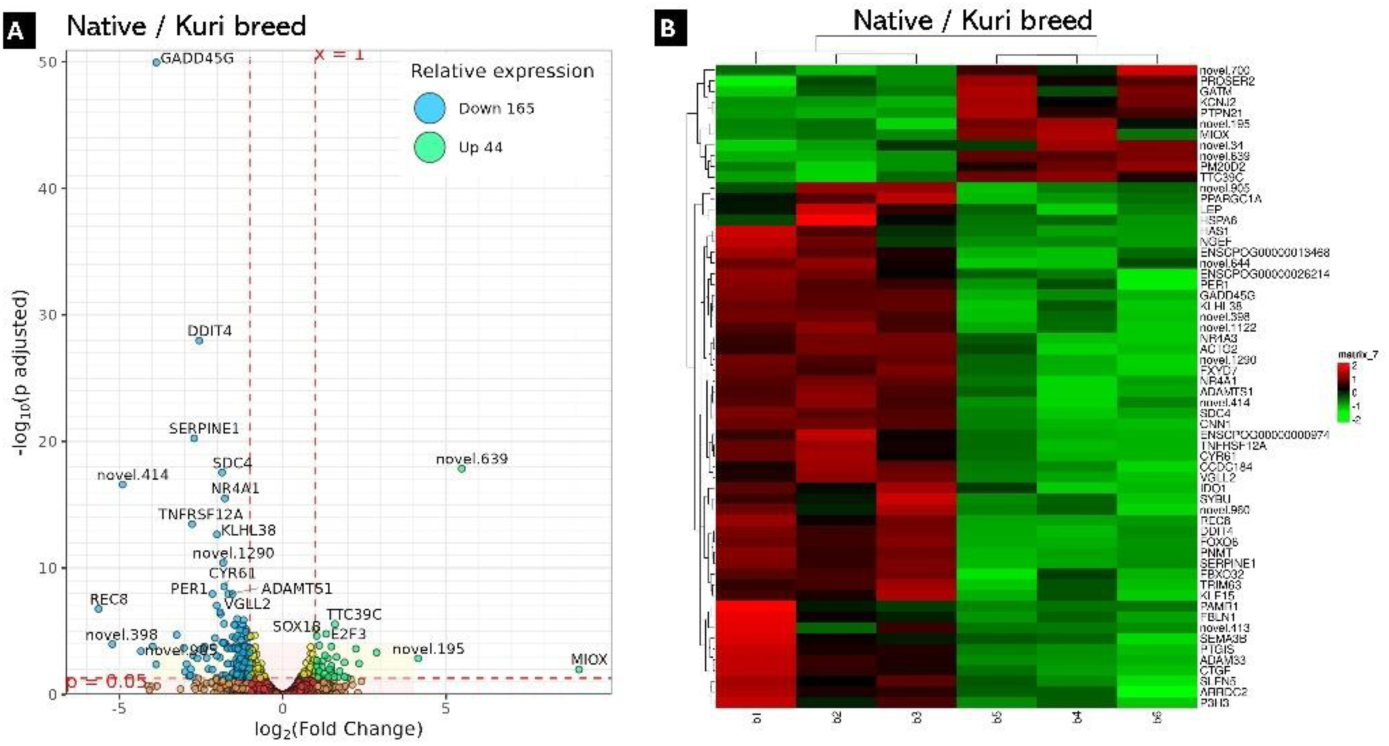
Differential expression analysis in global transcriptomic. (A) Volcano plot for differentially expressed of Native and Kuri breed guinea pig and (B) Heatmap profile of differential expression gene for native and Kuri breed guinea pig. Global color scale shows upregulated (red) and downregulated (green) expression levels.

### Enrichment analysis

The KEGG enrichment analysis showed that the differentially expressed genes (DEGs) in kuri breed, compared to native guinea pigs, were enriched in several metabolic pathways. These include the metabolism of glycine, serine, and threonine, arginine biosynthesis, and the metabolism of nicotinate, nicotinamide, ascorbate, aldarate, and carbohydrates, which are important for protein synthesis and energy availability. Signaling pathways such as the Notch pathway and viral carcinogenesis, involved in cell development, were also identified, along with hormonal pathways such as steroid hormone biosynthesis and ovarian steroidogenesis, which influence physiological processes (Fig 4A).

**Fig 4.**
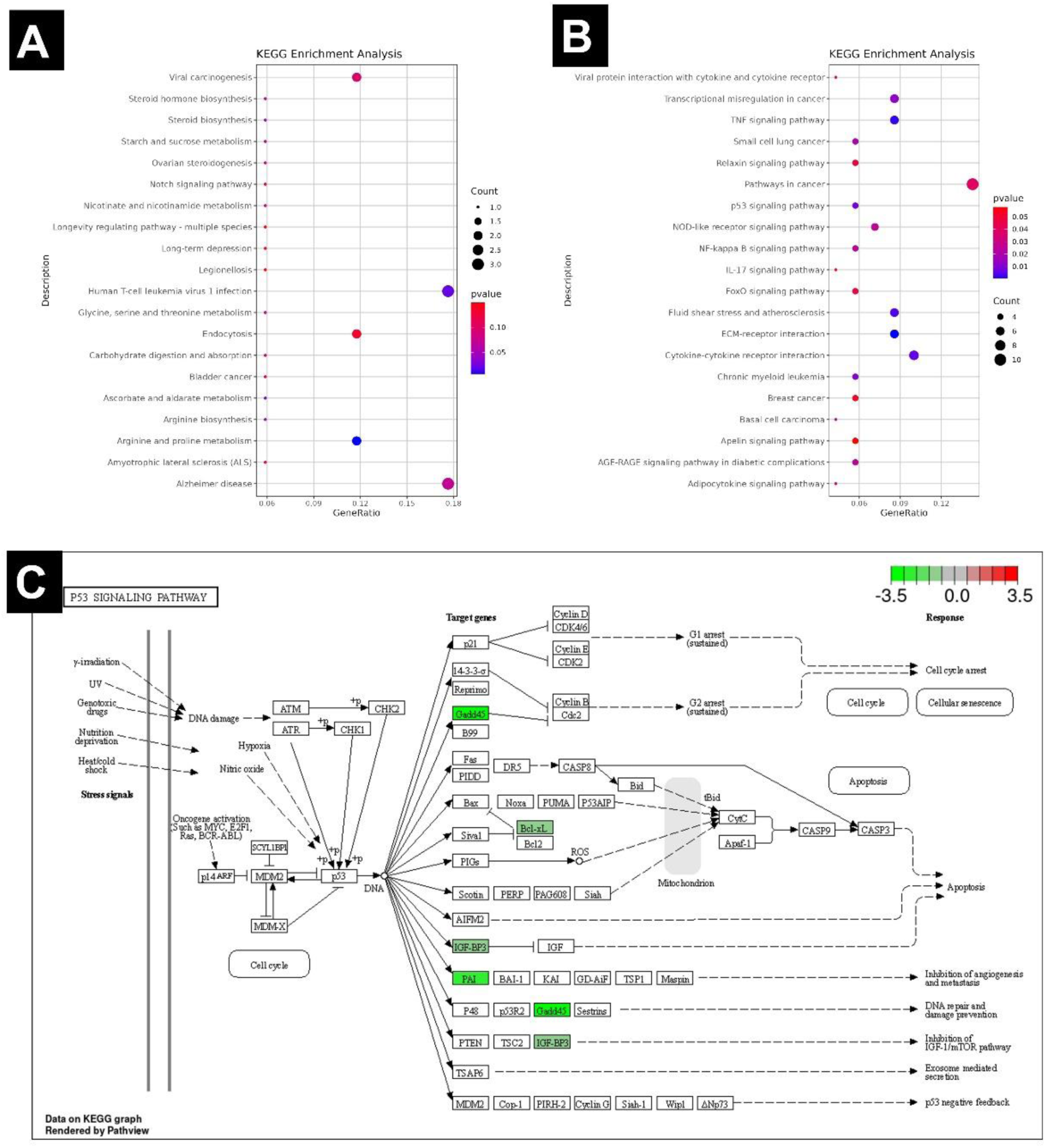
KEEG enrichment scatter plot for differential expression gene in native and Kuri breed guinea pig. (A) KEGG enrichment scatter plot for upregulated genes, (B) for downregulated genes and (C) P53 signaling pathway.

On the other hand, the KEGG analysis for underexpressed pathways revealed routes related to cellular stress and atrophy (Fig 4B). The FoxO signaling pathway, involved in the regulation of muscle atrophy through protein degradation [39], suggests a reduced activation of atrophy in these guinea pigs. Additionally, the p53 pathway, involved in apoptosis, DNA repair and damage prevention, inhibition of angiogenesis and metastasis, as well as inhibition of the IGF-1/mTOR pathway (Fig 4C), is also underexpressed, suggesting a lower activation of pro-apoptotic responses and inhibition of cell growth. In turn, reduced activity in the NF-kappa B pathways and cytokine-receptor interactions further supports a reduction in the inflammatory and stress response in guinea pigs with greater musculature.

### Identification and Annotation of Variants

Variants in the MSTN gene region were identified in the Kuri breed musculature guinea pigs, while no variants were detected in Native guinea pigs (Table 3). In sample B4, an INDEL corresponding to a frameshift deletion was detected, annotated as c.621delT. This mutation alters the reading frame, likely affecting the structure and function of the myostatin protein by producing a truncated protein. This type of mutation is classified as having HIGH impact.

**Table 3.**
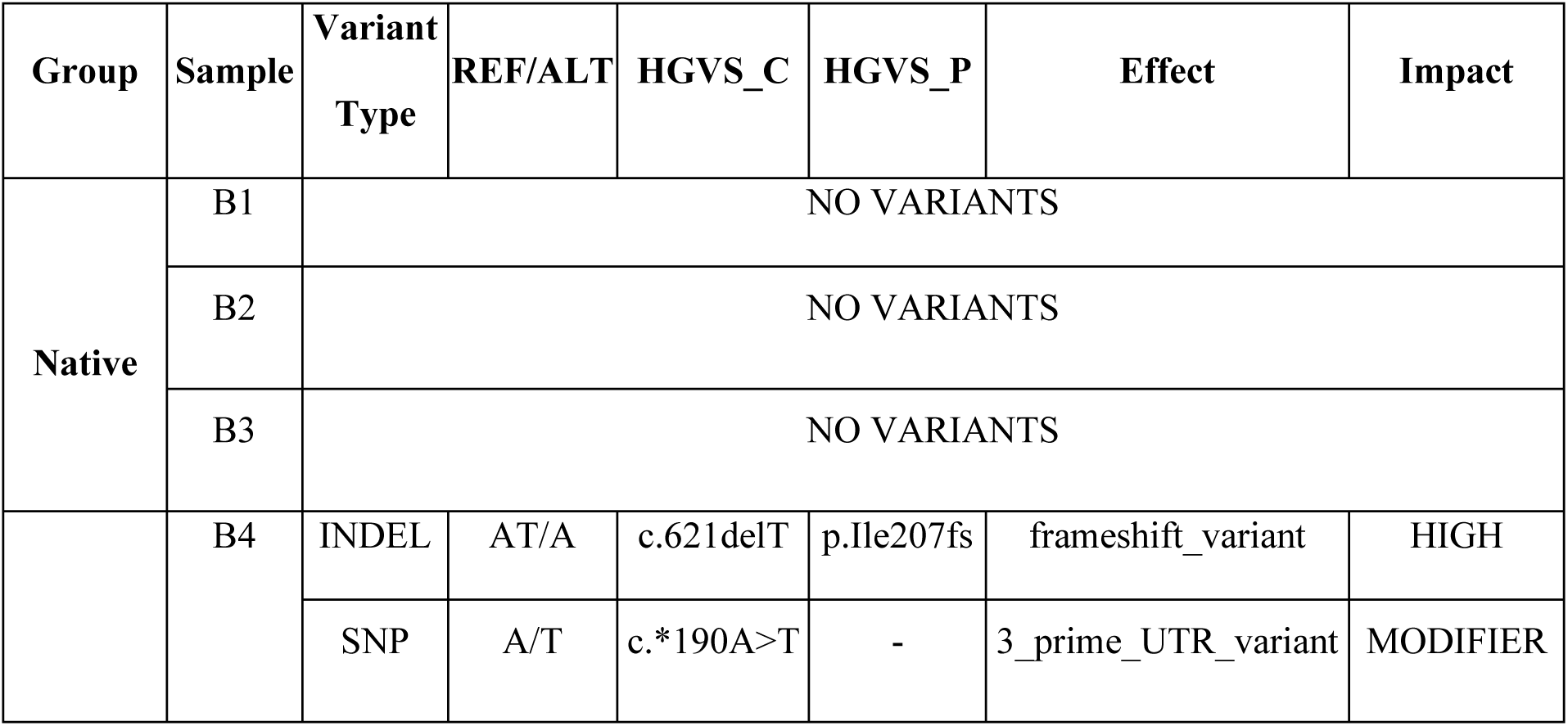
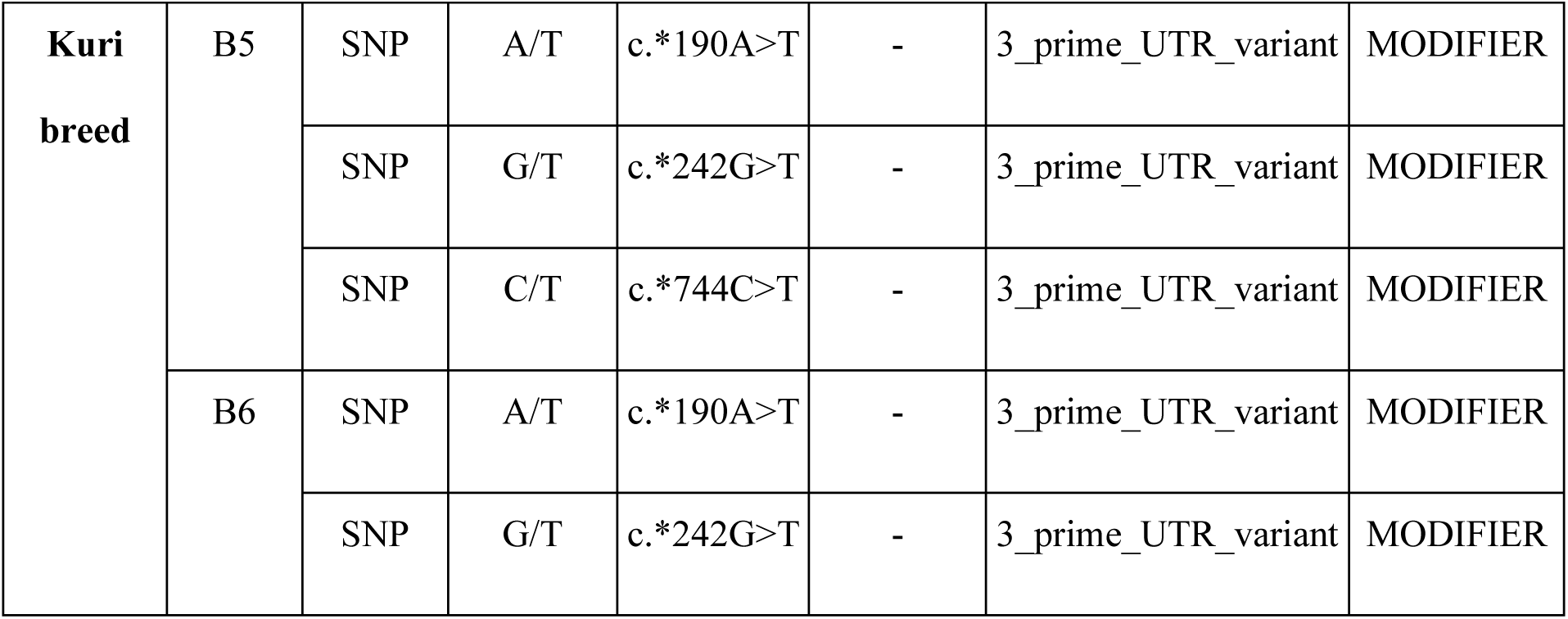
Genetic variants in the MSTN gene. Columns include the sample ID, type of variant, reference and alternate alleles (REF/ALT), HGVS_C annotation (cDNA level), HGVS_P annotation (protein level), predicted effect of the variant and the impact classification.

In samples B4, B5, and B6, a shared SNP was identified in the 3’ UTR region of the MSTN gene, annotated as *c.190A>T. This variant does not directly affect the protein sequence, but it is classified as a MODIFIER impact variant, suggesting that it may influence the post-transcriptional regulation of the gene, potentially affecting the stability or translation of the mRNA. In addition to the *SNP c.190A>T, in samples B5 and B6, other SNPs were found in the 3’ UTR region, specifically *c.242G>T and *c.744C>T, which were not present in B4.

## DISCUSSION

Our bulk RNA-seq analysis revealed significant differences in the expression of key genes related to muscle hypertrophy in kuri breed compared to native guinea pigs, suggesting a strong association with mutations in the MSTN gene and the activation of critical molecular pathways for muscle growth and degradation.

One of the most notable findings was that the MSTN gene showed slightly elevated expression in kuri breed guinea pigs, which contrasts with its known function as a muscle growth inhibitor. However, the variant analysis revealed that only these kuri breed guinea pigs presented mutations. Variants in the 3’ UTR region were identified, which may be involved in regulating MSTN expression, particularly the c.*190A>T variant, which was found in all three samples. Previous studies have shown that mutations in the 3’ UTR region can block MSTN translation, resulting in “double-muscle” phenotypes [40–42]. Additionally, a frameshift variant was found that causes a reading frame shift, likely resulting in a truncated or non-functional protein. This type of mutation, which alters the reading frame, has been associated with phenotypes characterized by high carcass yield, lower fat grade [43,44], increased musculature [45], and hyperplasia [46]. The absence of mutations in native guinea pigs suggests that MSTN mutations are linked to the enhanced muscle phenotype in kuri breed guinea pigs. Furthermore, the slight overexpression of MSTN in these guinea pigs may represent a compensatory mechanism by the organism to attempt to regulate excessive muscle growth. However, due to the mutations present, this overexpression does not seem to have the expected effect of inhibiting muscle growth.

In addition to MSTN, several metabolic and signaling pathways were differentially regulated between kuri breed and native guinea pigs. One of these is the ubiquitin-proteasome pathway, associated with muscle protein loss through accelerated protein degradation [47–49], while its inhibition reduces atrophy in mice [50]. Specifically, we found two underexpressed ubiquitin ligases in kuri breed guinea pigs: Tim63 (also known as MuRF1) and Fbxo32 (MAFbx), which have been shown to significantly contribute to muscle atrophy [47,51,52], and their inhibition prevents it [53,54], so this inhibition found could be protecting the skeletal muscle of kuri breed guinea pigs from protein degradation. Furthermore, Fbxo32 expression has been found to be mediated by transcription factors FoxO and TNF-α [39,55], while MuRF1 is activated by the NF-KB pathway [56,57], which is also influenced by TNF-α in skeletal muscle, suggesting that the underexpression of these pathways (Fig 4B) is also associated with muscle protection in kuri breed guinea pigs.

Another relevant gene is Ddit4, which is underexpressed in kuri breed guinea pigs. This gene, elevated in skeletal muscle under atrophic conditions caused by food deprivation, cancer cachexia, spinal muscular atrophy, drugs, among others [58–62], inhibits the mTOR pathway [60,63–65], which controls muscle protein synthesis [66–68]. Ddit4 underexpression could be allowing greater activation of protein synthesis in kuri breed guinea pigs, promoting muscle growth. Additionally, the underexpression of the p53 signaling pathway, which inhibits mTOR through the IGFBP3 protein [69–71], supports the hypothesis of increased mTOR activity linked to muscle hypertrophy. The p53 pathway in skeletal muscle has been associated with muscle atrophy due to cachexia [72] and senescence [72,73]. Finally, the overexpression of ZNF383, which has been observed to repress the p53 pathway [74], could also be contributing to the reduced atrophy observed.

Regarding the function of the GADD45G gene, we observed that it is underexpressed in synthetic kuri breed guinea pigs. This gene encodes for growth arrest and DNA damage-inducible gamma protein [75]. It has been attributed with a role as a growth inhibitor and apoptosis inducer, as its expression has been shown to suppress tumor cell growth [76,77], acting as an inhibitor of cdc2/cyclinB1 kinase at the S and G2-M cell cycle checkpoints [78]. Additionally, GADD45a, another protein in the same family, has been identified as a critical mediator of atrophy [79–81] through myonuclear remodeling. However, we did not find significant changes in its expression in our study, only in GADD45g, which may indicate that this protein is substituting for its homolog in the regulation of atrophy. Additionally, the GADD45 family has been reported to be involved in reactive oxygen species (ROS) production, as Gadd45-deficient mice were observed to have decreased ROS production [82].

Another downregulated protein is SERPINE1, which acts as an inhibitor of urokinase plasminogen activator (uPA), resulting in inhibition of extracellular matrix degradation [83], which is important in muscle regeneration [84] and exercise-induced adaptation [85] because it allows the removal of damaged components, preparing the environment for the formation of new muscle fibers, releasing growth factors and cytokines that stimulate the proliferation and differentiation of myogenic cells [86]. Additionally, extracellular matrix restructuring has been shown to be important for angiogenesis in mouse models after physical activity was induced [87]. Other ECM-related genes found to be downregulated include CYR61, a protein associated with muscle atrophy that participates in fiber-type switching during sarcopenia [88] and reduces the proliferation of muscle progenitor cells in skeletal muscle during aging [89]. We also found SDC4 to be upregulated in Native guinea pigs, where overexpression of syndecan-4 has been reported to inhibit myogenesis, showing a smaller total myotube area [90], suggesting that avoiding the overexpression of this gene in kuri breed guinea pigs has supported their muscle hypertrophy.

On the other hand, the overexpression of genes related to energy metabolism and angiogenesis was observed. Among them is GATM, an important enzyme in creatine biosynthesis [91]. In mammals, most creatine comes from its synthesis in the liver and kidneys [91,92], later being taken up by tissues through creatine transport proteins [93]. However, a study demonstrated that by blocking the creatine transporter SLC6A8, GATM levels and the rate of creatine biosynthesis in skeletal muscle increased [94]. This could suggest that during muscle hypertrophy, there is a higher demand for creatine, which cannot be satisfied solely through extracellular creatine, making it necessary for GATM to increase its synthesis in skeletal muscle. Similarly, the overexpression of MIOX, which encodes the myo-inositol oxygenase enzyme that catabolizes myo-inositol to D-glucuronate [95], entering the pentose phosphate pathway [96], may play a role in the muscle’s adaptation to increased energy demands in kuri breed guinea pigs, as its expression in horses has been observed to contribute to muscle adaptation during endurance exercise [97], and it has been found to be downregulated in Duchenne muscular dystrophy models in dogs [98].

The overexpression of ANGPT1 was also identified, which binds to Tie-2 receptors, activating pathways for the formation of new blood vessels in muscle [99]. In the case of kuri hibrid guinea pigs, angiogenesis plays a crucial role, as fiber hypertrophy leads to the accumulation of muscle mass, and angiogenesis becomes essential for supplying oxygen and nutrients and removing metabolic waste [100]. Microcirculation must be sufficient to meet the metabolic demands imposed on muscle [101]. This protein has also been involved in myogenesis, as it stimulates the proliferation and differentiation of muscle stem cells into myotubes, while its inactivation has been shown to decrease these processes [99,102,103]. Finally, the protein TC39C was found to be upregulated, where it has been observed that in the skeletal muscle of mice, it reaches maximum expression between late proliferation and early differentiation [104]. Additionally, this protein has been found to be enriched in a new hybrid goat breed with superior growth characteristics [105].

Together, these findings reinforce the idea that MSTN mutations, along with the differential regulation of several key molecular pathways, are modulating the enhanced muscle phenotype in kuri breed guinea pigs. However, future studies investigating the functional role of these variants and pathways will be necessary to fully understand the molecular mechanisms behind these phenotypes.

## CONCLUSSION

This comparative transcriptomic analysis between native and kuri breed guinea pigs revealed that mutations in the *MSTN* gene, including variants in the 3’ UTR region and a frameshift mutation, play a key role in regulating muscle growth by disrupting its inhibitory function. These mutations, along with the downregulation of pathways related to muscle degradation and the upregulation of genes associated with energy metabolism and angiogenesis, explain the muscle hypertrophy observed in the kuri breed guinea pigs. This study provides a first insight into how this breed has developed genetic mechanisms that promote muscle hypertrophy, paving the way for future research and its potential impact on guinea pig production.

## MATERIALS AND METHODS

The study was carried out following the Ethics regulation of scientific research of Universidad Nacional Agraria La Molina (CEI-UNALM-006/2022).

### Sample Collection

The male guinea pigs belong to the Experimental Center, La Molina of the National Institute of Agrarian Research (INIA, Lima, Peru). They were raised in a controlled environment and fed appropriate food for their age. Three native and three Kuri breed males were randomly selected with an average live weight at birth (107 and 155 g) and eight weeks (535 and 791 g), respectively. The animals were sacrificed at 78 days of age under standardized procedures in a guinea pig commercial slaughterhouse and then muscle tissue samples were collected from the hind leg. The samples were immediately frozen at-80 °C until processing.

### Muscle histology

The samples were fixed in 4% paraformaldehyde for 24 hours at 4°C. After that time, they were successively dehydrated for 1 hour in ethanol gradients of 80%, 90% and 3 times in 100%. They were then incubated 3 times in Xylol and then embedded in paraffin and sectioned into 10 µm thick sections and each tenth section of the muscle tissue was mounted on slides. Subsequently, they were stained with hematoxylin and eosin. The images were captured using a histological microscope (Zeiss®, PrimoStar, Germany) attached with a digital camera at X40 magnification.

### RNA Preparation and Sequencing

Total RNA of each muscle sample was extracted from approximately 50 mg of frozen tissue using Direct-zol RNA Miniprep - Zymo Research (Zymo Research, CA, USA) following the manufacturer’s instructions. The quality of the RNA samples was checked using an Agilent 2100 Bioanalyzer (Agilent Technologies, CA, USA). The total RNA of 6 samples were used for sequencing with UltraTM RNA Library Prep kit for Illumina® (NEB, MA, USA); all the standards and procedures were performed following the manufacturer’s protocols. After the quality control using Agilent 2100 Bioanalyzer and ABI StepOnePlus Real-Time PCR System (ABI, Vernon, CA, USA), the library preparations were sequenced on an Illumina Hiseq 4000 platform (Illumina, Novogene, CA, USA) and 100 bp paired-end reads were generated. The RNA library construction and RNA-sequencing services were provided by Novogene USA (Fig 5).

**Fig 5.**
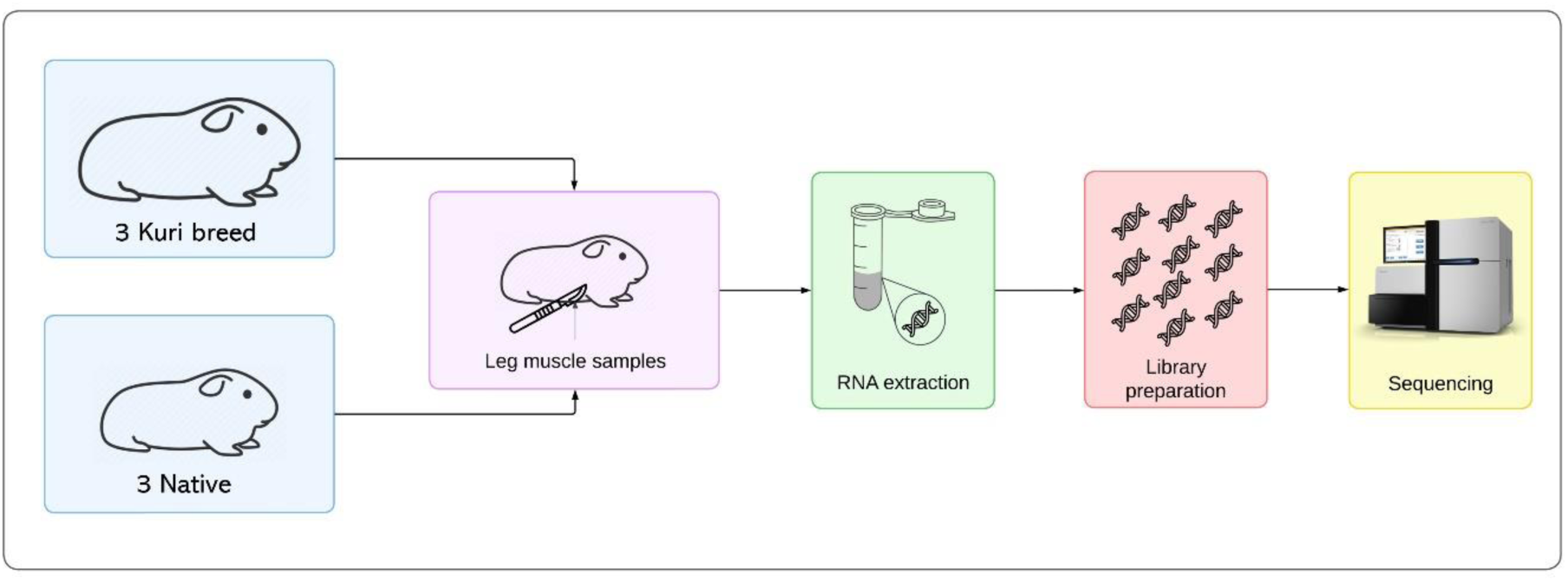
Workflow for RNA extraction and library preparation for muscle sample sequencing

### RNA-Seq Data Analysis

Clean reads were generated by removing those containing adapters, poly-N sequences, or exhibiting low quality from the unprocessed reads using Fastp software [24]. These clean reads were aligned to the *Cavia porcellus* reference genome from Ensembl (GCA_000151735.1) using HiSAT v2.0.5 [25]. Subsequently, assemblies of the aligned reads were performed using StringTie v1.3.3b [26]. Then, the assembled fragments were compared to the reference transcriptome to discover new genes. The quantification of gene expression was performed utilizing Salmon software [27], with the Ensembl *Cavia porcellus* transcriptome as reference, including the new genes identified in the previous step. All bioinformatics analyses were conducted on the O-HPC at the Universidad Nacional Agraria La Molina

### Differential Expression Analysis

Differential expression analysis between the two groups was performed using the DESeq2 package v1.40.2 [28], followed by p-value correction using the Benjamini-Hochberg method [29]. Genes that met the criteria of p < 0.05 and absolute Fold Change ≥ 1 were considered differentially expressed. These genes were used to create a heatmap with normalized and standardized (z-score) counts. All analyses, including filtering, dimensionality reduction, and generation of other plots, were carried out in R software.

### Pathway Enrichment Analysis

Enrichment analysis was performed using the differentially expressed genes. For pathway enrichment analysis, the Kyoto Encyclopedia of Genes and Genomes (KEGG) database was used [30]. The significance of GO and KEGG terms was assessed using the clusterProfiler package v4.8.2[31], which includes correction for gene length bias. GO and KEGG terms with corrected p-values < 0.05 were considered significantly enriched by the differentially expressed genes.

### Identification and Annotation of Variants

Variant analysis was performed using GATK v4.1.1.0 [32] which allowed for the detection of SNPs and indels from the aligned reads. Subsequently, the variants were filtered using GATK’s Variant Filtration module, applying quality criteria such as QD (Qual by Depth) < 2.0, FS (Fisher Strand Bias) > 30.0 to remove variants with strand bias, and DP (Depth) < 10 to exclude those with low read coverage. The filtered variants were then annotated with SnpEff v4.3 [33], allowing for the identification and classification of variant types and their impacts, ranging from high to modifier, to assess their biological relevance.

## Acknowledgement

This research was supported by PROCIENCIA, Peru Grant PROCIENCIA 417-2019-FONDECYT.

## Author contributions

Conceptualization, EA, EM; Data curation, SC, EM; Formal analysis, SC, EM; Funding acquisition, EA, EM, LC; Investigation, EM, CY, LC; Methodology, EM, CY, SC; Project administration, EA, EM; Software, SC, EM; Supervision, EA, EM; Validation, CY, EM; Visualization, SC, EM; Writing – original draft preparation, SC, EM; Writing – review & editing, SC, LC, EM.

## SUPPLEMENTARY MATERIAL

Fig S1. Histology of the muscle and live weight of animals used in the experiment. A) Histological section of the guinea pig muscle stained with Hematoxylin-eosin (40x). B) Evolution of live weights from birth to 8 weeks of age of native and kuri breed guinea pigs. BW: Birth weight, WW: weaning weight; W4w: Weight at 4 weeks; W6w: weight at 6 weeks and W8w: Weight at 8 weeks.

Table S1. Summary of gene differential expression analysis between kuri breed and native guinea pigs, Up expression.

Table S1. Summary of gene differential expression analysis between kuri breed and native guinea pigs, Down expression.

